# Susceptibility Patterns of Multiple Antibiotic-Resistant Bacteria from Wound and Urine Samples to the Extract of *Spondias Mombin* (Linn)

**DOI:** 10.1101/2024.06.26.600810

**Authors:** AF Okiti, MK Oladunmoye, AO Ogundare

## Abstract

This study evaluates the antibacterial activity of *Spondias mombin* L. against multiple antibiotic-resistant bacteria isolated from wound and urine samples of patients attending five (5) selected hospitals in Akure, Ondo State, Nigeria. A total of 313 bacterial isolates were isolated from 353 samples of wound and urine using standard bacteriological procedures, with *Pseudomonas aeruginosa* and *Staphylococcus aureus* being the most predominant in urine and wound samples, respectively. The methanolic extract of *S. mombin* was the most effective against wound isolates, while the aqueous extract was the most effective against urine isolates. The results showed that the methanol extract of *S. mombin* had a zone of inhibition of 24.00±0.00, 30.67±0.33 and 19.33±0.33 mm respectively, against *S. aureus, S. epidermidis* and *P. aeruginosa* at 100 mg/ml. The aqueous extract had a zone of inhibition of 24.67±0.33, 27.33±0.33, 18.67±0.33, 24.67±0.33, 23.67±0.33, 21.33±0.33 and 21.67±0.33 mm against *Escherichia coli, Klebsiella pneumoniae, P. aeruginosa, Proteus mirabilis, S. aureus, S. saprophyticus* and *Trichomonas vaginalis* respectively, at 100 mg/ml. The phytochemical constituents of the extracts include alkaloids, anthraquinones, cardiac glycosides, flavonoids, phenols, saponins, steroids and tannins. These compounds may be responsible for the antibacterial activity of *S. mombin* against the multiple antibiotic-resistant bacterial isolates. The findings of this study demonstrates the potential of *S. mombin* as an alternative treatment for multiple antibiotic-resistant bacteria from wound and urine.

## Introduction

Antibiotic resistance is a major global health crisis that threatens the effective treatment of infectious diseases. When bacteria acquire the ability to withstand antimicrobial medications that are meant to kill them, antibiotic resistance arises. This resistance can lead to the emergence of “superbugs” that are difficult or impossible to treat with currently available antibiotics [1]. The overuse and inappropriate use of antibiotics are major drivers of the emergence and spread of antibiotic resistance [2]. In addition, the widespread use of antibiotics in agriculture and aquaculture has also contributed to the development of antibiotic resistance in bacteria [3]. The consequences of antibiotic resistance are significant and potentially catastrophic. Antibiotic resistance causes 700,000 global deaths annually, with rising numbers expected unless urgent action is taken. Economic costs include longer hospital stays, expensive treatments, and decreased productivity [4]. In the face of this crisis, there is a pressing need to identify alternative antimicrobial agents that can effectively treat multiple drug-resistant infections.

Plant-derived compounds have been used for centuries in traditional medicine for their antimicrobial properties [5]. Plant-based antimicrobials offer a promising alternative to synthetic ones, potentially reducing antibiotic resistance, attracting increasing scientific interest in addressing the problem [6]. One such plant is *Spondias mombin*, commonly known as the yellow mombin or hog plum in English, known has Iyeye in Yoruba, Ijikara (Igbo), Tsardar masar (Hausa). *S. mombin*, originating from Central and South America, is a fruit-bearing tree that thrives in tropical regions. It has a long history of use in traditional medicine for the treatment of a variety of infections [7]. The fruit, leaves and bark of the tree have been used to treat a range of conditions, including stomachache, diarrhea, wound, fever and urinary tract infections [7].

Recent studies highlight the potential of *S. mombin* as an antimicrobial agent due to its phytochemicals, particularly saponins, which exhibit potent antimicrobial activity against various bacteria and fungi [7]. *S. mombin* fruit, known for its antimicrobial and anti-inflammatory properties, has hypoglycemic effects in animal studies, suggesting potential as a natural diabetes treatment [8]. *S. mombin* leaf extracts have also been shown to have hypolipidemic effects in animals, meaning that they may help to lower levels of cholesterol and other lipids in the blood [8]. Despite the various medicinal properties of *S. mombin*, more research is needed to fully understand its therapeutic potential and to identify the active components responsible for its effects.

## Materials and Methods

### Study Population

Seventy-one wound samples and 282 urine samples were collected from patients in five Akure hospitals, including diabetic, accident, burns, and postoperative wounds, and from patients with and without urinary tract infection history.

### Wound and Urine Sample Collection

Wounds were cleansed with sterile normal saline, and wound swabs were collected from all participants using sterile moistened cotton swabs in an aseptic manner. The swabs were then placed in an icepack container and transported to the laboratory [9]. Urine samples were collected using sterile universal bottles and transferred to the research laboratory for further processing [10].

### Isolation and Identification Method

Wound swab samples were immediately placed in Mueller Hilton broth, followed by streaking on Mannitol Salt Agar (MSA), Nutrient Agar, and Blood agar plates. The plates were incubated aerobically at 37°C for 24 hours. Presumptive identification of bacteria was based on their cultural characteristics on each agar plate. Representative colonies were sub-cultured on MSA plates and Nutrient agar, and further incubated at 37°C for 24 hours. The distinct, well-isolated colonies were then studied for their cultural and morphological characteristics. Gram staining and biochemical tests were conducted to confirm the identification of the isolates [11].

For urine samples, 1 μl of urine was spread quantitatively on MacConkey agar, and CLED agar, and incubated aerobically at 35°C for 24 hours. Similar procedures for presumptive identification, sub-culturing, incubation, and confirmation were followed for the urine samples [10]. Preliminary characterization of the isolates was conducted using Gram staining, morphological examination, cultural characteristics and biochemical tests according to Shoaib *et al*. [12].

### Antibiotics Susceptibility Testing

The susceptibility of the isolates to antibiotics was determined using the disk diffusion method as described by Cheesbrough [13]. Gram-positive isolates were tested against eight commercially available antibiotics CAZ(30µg), CRX(30µg), GEN(10µg), CTR(30µg), ERY(5µg), CXC(5µg), OFL(5µg), AUG(30µg), while Gram-negative isolates were tested against eight different antibiotics CAZ(30µg), CRX(30µg), GEN(10µg), CXM(10µg), OFL(5µg), AUG(30µg), NIT(30µg), CPR(10µg). The zones of inhibition were compared with the standard guidelines.

### In-vitro Antibacterial Studies

The leaves were dried and ground into coarse powder. Extraction and standardization of plant extracts were carried out following established procedures as described by Okiti and Osuntokun [14]. The agar well diffusion method was employed for the antibacterial studies. The dried plant extract was reconstituted with sterile distilled water and ethyl acetate to obtain different concentrations of 100mg/ml, 50mg/ml, 25mg/ml and 12.5mg/ml. Bacterial strains were cultured and spread on Mueller-Hinton agar plates, and wells were made in the agar. The plant extracts were introduced into the wells at different concentrations, and ciprotab 2mg/ml was used as control. The plates were incubated at 37^0^C and the zones of inhibition were measured to determine the antibacterial activity of the plant extract [15].

### Determination of Minimum Inhibitory Concentration (MIC) and Minimum Bactericidal Concentration (MBC)

The MIC of the extracts against the test organisms was determined using the broth dilution method described by Rankin and Coyle [16]. Serial dilutions of the extracts were prepared to obtain extract concentrations of 100mg/ml, 50mg/ml, 25mg/ml and 12.5mg/ml, and their inhibitory concentrations were recorded. The MBC was determined using the method established by the National Committee for Clinical Laboratory Standard [17]. Samples that did not show visible growth after incubation were streaked on Nutrient Agar plates to determine the minimum concentration of the extract required to kill the organisms. The lowest concentration indicating a bactericidal effect was recorded as the MBC. The qualitative phytochemical tests were carried out using the method described by the following authors: Alkaloid test [18], Anthraquinone test [19, 20], Cardiac Glycosides test [18, 20]. Flavonoid test [18, 19, 21], Phenol test [19], Saponin test [18, 22], Steroid test [18], Tannin test [19, 22].

### Statistical Analysis

Data obtained were subjected to one way analysis of variance (ANOVA) and Duncan’s New Multiple Range Test at 95% confidence level using SPSS 20.0 version. Differences were considered significant at P ≤ 0.05.

## RESULTS

A total of 353 samples were analyzed for the presence of multiple antibiotic-resistant bacteria. 59 bacterial isolates were recovered from 67 wound samples collected. The predominant bacteria isolated from the infected wounds were *S. aureus* 39 (66.10%) followed by *P. aeruginosa* 15 (25.42%), and *S. epidermidis* 5 (8.47%). A total of 254 bacterial isolates were recovered from 282 urine samples. The predominant bacteria isolated from the urine samples were *P. aeruginosa* 84 (33.07%), followed by *S. aureus* 71 (27.95%), *S. saprophyticus* 41 (16.14%), *E. coli* 25 (9.84%), *T. vaginalis* 19 (7.48%), *K. pneumonia* 9 (3.54%), and *Proteus* sp. 5 (1.97%).

## DISCUSSION

This study investigated the susceptibility patterns of multiple antibiotic-resistant bacteria isolated from wound and urine samples to the extract of *S. mombin*. The findings revealed a high prevalence of multiple antibiotic-resistant bacteria in both sample types, with *P. aeruginosa* being the most commonly isolated bacterium in urine samples and *S. aureus* being the most commonly isolated bacterium in wound samples. These results align with previous research indicating the prominence of *S. aureus* and *P. aeruginosa* as common antibiotic-resistant bacteria. Liu and Qin [23] reported similar findings, identifying these two bacteria as among the top five antibiotic-resistant pathogens in recent years, reflecting their well-known association with nosocomial infections and their ability to develop antibiotic resistance. The high prevalence of antibiotic-resistant bacteria in clinical samples highlights the need for effective antimicrobial strategies. The identification of other bacterial species in wound and urine infections highlights the complexity of the microbial landscape, guiding the development of appropriate therapeutic approaches.

Table 1 and Table 2 present the resistance profiles of bacterial isolates, revealing that *S. aureus* and *P. aeruginosa* showed the highest MAR indices of 0.8, indicating resistance to 80% of tested antibiotics. A strain of *S. epidermidis* displayed susceptibility to all antibiotics, indicating a generally less resistant bacterial strain compared to other *Staphylococcus* species. These results align with previous studies that have identified *S. aureus* and *P. aeruginosa* as major contributors to antibiotic resistance [23]. Table 2 provides further insights into the antibiotic susceptibility profiles of bacteria isolated from urine samples. The results indicate that *K. pneumoniae* exhibited the highest MAR index of 0.7, reflecting resistance to 70% of the antibiotics tested. This finding is in line with previous research that has highlighted the high prevalence of multidrug-resistant *K. pneumoniae* strains [24]. Similarly, *P. aeruginosa* showed resistance to 60% and *Proteus mirabilis* had a lower resistance index, this finding does not agree with existing studies that have recognized *Proteus* mirabilis as a high commonly encountered multidrug-resistant pathogen [25].

**Table 1:** Antibiotic sensitivity profile of bacteria isolated from wound.

| Isolates | Antibiotics to which isolates were resistant | MAR Index |
| --- | --- | --- |
| *S. aureus* | CAZ, CRX, GEN, CTR, ERY, CXC, OFL, AUG | 0.8 |
| *P. aeruginosa* | CAZ, CRX, GEN, CXM, OFL, AUG, NIT, CPR | 0.8 |
| *P. aeruginosa* | CAZ, CRX, CXM, OFL, AUG, CPR | 0.6 |
| *S. aureus* | CAZ, CRX, CXC, AUG | 0.4 |
| *S. epidermidis* | - | 0 |
| *S. epidermidis* | CAZ | 0.1 |
| *S. aureus* | GEN, OFL | 0.2 |

**Key:** CAZ – Ceftazidime (30µg), CRX - Cefuroxime (30µg), GEN - Gentamycin (10µg), CXM – Cefixime (10µg), OFL – Ofloxacin (5µg), AUG – Augmentin (30µg), NIT – Nitrofurantoin (30µg), CPR – Cefpiroma (10µg).

**Table 2:** Antibiotic sensitivity profile of bacteria isolated from urine.

| Isolates | Antibiotics to which isolates were resistant | MAR Index |
| --- | --- | --- |
| *K. pneumoniae* | - | 0 |
| *P. mirabilis* | CRX, OFL, AUG, CPR | 0.4 |
| *E. coli* | AUG | 0.1 |
| *P. aeruginosa* | CAZ, CRX, GEN, CXM, AUG, NIT | 0.6 |
| *S. saprophyticus* | CAZ, CRX, CTR, CXC, AUG | 0.5 |
| *K. pneumoniae* | CAZ, CRX, GEN, CTR, ERY, CXC, AUG | 0.7 |
| *S. aureus* | CAZ, CRX, ERY, CXC, AUG | 0.5 |
| *P. mirabilis* | AUG | 0.1 |
| *S. aureus* | CAZ, CRX, CTR, CXC, AUG | 0.5 |

**Key:** CAZ – Ceftazidime (30µg), CRX - Cefuroxime (30µg), GEN - Gentamycin (10µg), CXM – Cefixime (10µg), OFL – Ofloxacin (5µg), AUG – Augmentin (30µg), NIT – Nitrofurantoin (30µg), CPR – Cefpiroma (10µg).

From table 3, 4, and 5, the aqueous extract of *S. mombin* showed susceptibility against *S. aureus, S. epidermidis*, and *P. aeruginosa*, with zone diameters greater than those produced by ciprotab (24.67±0.33d, 28.33±0.33d), suggesting that the aqueous extract of *S. mombin* may possess stronger antimicrobial activity against these bacterial isolates. The methanolic extract also showed susceptibility to these bacteria, with larger zones of inhibition than the positive control. The n-Hexane extract showed varying degrees of effectiveness against antibiotic-resistant isolates from wound samples, with zone diameters varying from 0.00±0.33a to 8.33±0.33a.

**Table 3:**
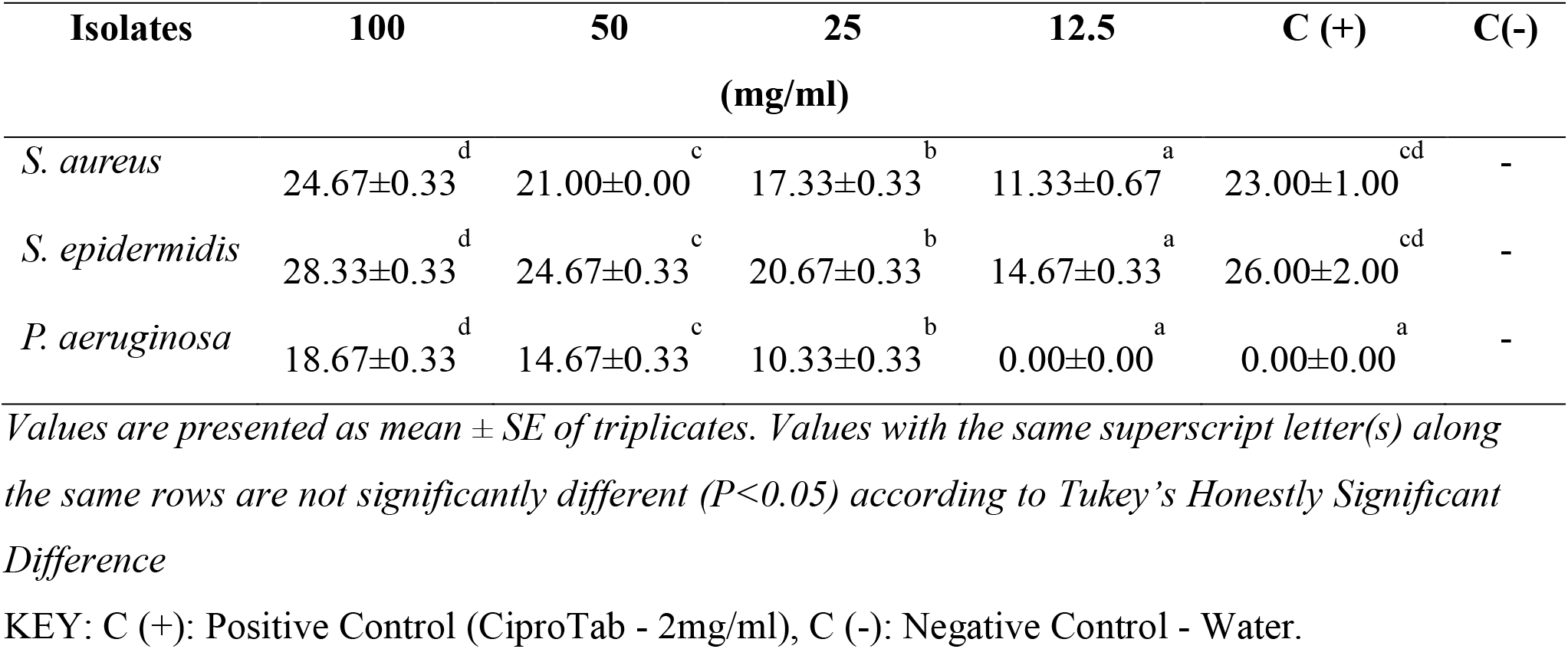
Antibacterial activity of aqueous extract of on isolates from wound samples.

| Isolates | 100 | 50 | 25 | 12.5 | C (+) | C(-) |
| --- | --- | --- | --- | --- | --- | --- |
|  | (mg/ml) |  |  |  |  |  |
| *S. aureus* | 24.67±0.33<sup>d</sup> | 21.00±0.00<sup>c</sup> | 17.33±0.33<sup>b</sup> | 11.33±0.67<sup>a</sup> | 23.00±1.00<sup>cd</sup> | - |
| *S. epidermidis* | 28.33±0.33<sup>d</sup> | 24.67±0.33<sup>c</sup> | 20.67±0.33<sup>b</sup> | 14.67±0.33<sup>a</sup> | 26.00±2.00<sup>cd</sup> | - |
| *P. aeruginosa* | 18.67±0.33<sup>d</sup> | 14.67±0.33<sup>c</sup> | 10.33±0.33<sup>b</sup> | 0.00±0.00<sup>a</sup> | 0.00±0.00<sup>a</sup> | - |

Values are presented as mean ± SE of triplicates. Values with the same superscript letter(s) along the same rows are not significantly different ( $P<0.05$ ) according to Tukey's Honestly Significant Difference
KEY: C (+): Positive Control (CiproTab - 2mg/ml), C (-): Negative Control - Water.

**Table 4:** Antibacterial activity of methanol extract on isolates from wound samples.

| Isolates | 100 | 50 | 25 | 12.5 | C(+) | C(-) |
| --- | --- | --- | --- | --- | --- | --- |
|  | (mg/ml) |  |  |  |  |  |
| *S. aureus* | 24.00±0.00<sup>d</sup> | 21.00±0.58<sup>c</sup> | 18.33±0.67<sup>b</sup> | 14.67±0.33<sup>a</sup> | 23.00±1.00<sup>cd</sup> | - |
| *S. epidermidis* | 30.67±0.33<sup>d</sup> | 26.67±0.33<sup>c</sup> | 22.67±0.33<sup>b</sup> | 18.00±0.00<sup>a</sup> | 26.00±2.00<sup>c</sup> | - |
| *P. aeruginosa* | 19.33±0.33<sup>d</sup> | 16.00±0.00<sup>c</sup> | 13.00±0.00<sup>b</sup> | 0.00±0.00<sup>a</sup> | 0.00±0.00<sup>a</sup> | - |

Values are presented as mean ± SE of triplicates. Values with the same superscript letter(s) along the same rows are not significantly different ( $P<0.05$ ) according to Tukey's Honestly Significant Difference
KEY: C (+): Positive Control (CiproTab - 2mg/ml), C (-): Negative Control - Methanol.

**Table 5:**
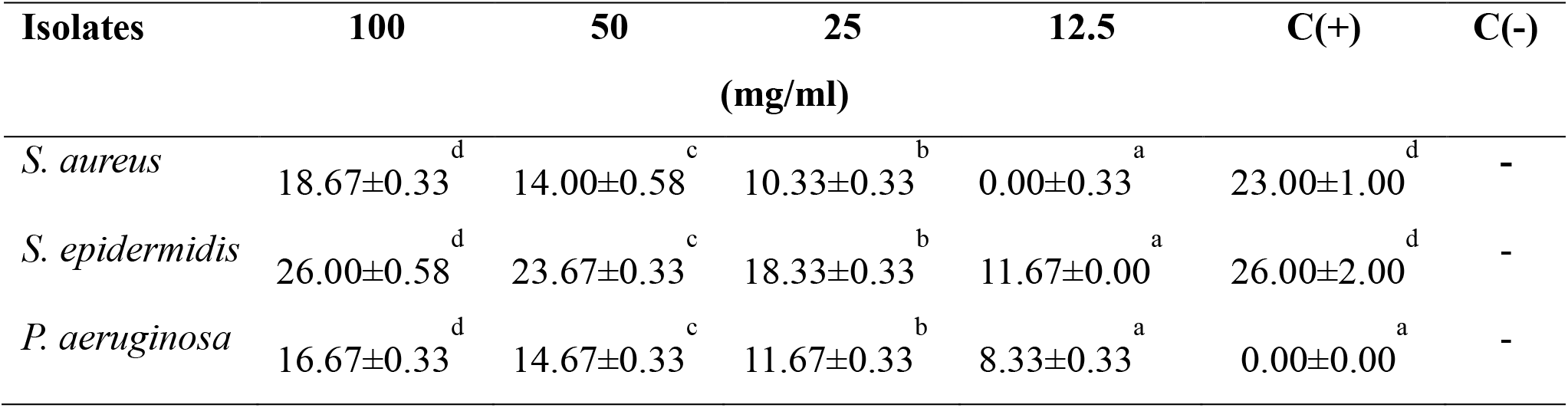
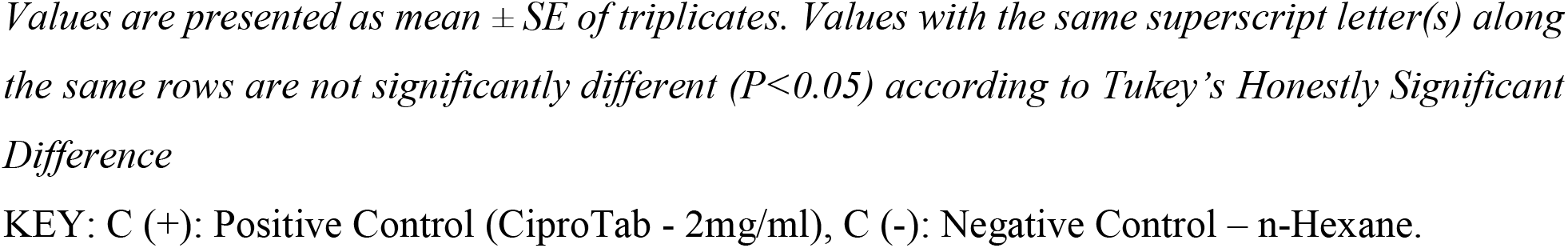
Antibacterial activity of n-Hexane extract on isolates from wound samples.

| Isolates | 100 | 50 | 25 | 12.5 | C(+) | C(-) |
| --- | --- | --- | --- | --- | --- | --- |
|  | (mg/ml) |  |  |  |  |  |
| *S. aureus* | 18.67±0.33<sup>d</sup> | 14.00±0.58<sup>c</sup> | 10.33±0.33<sup>b</sup> | 0.00±0.33<sup>a</sup> | 23.00±1.00<sup>d</sup> | - |
| *S. epidermidis* | 26.00±0.58<sup>d</sup> | 23.67±0.33<sup>c</sup> | 18.33±0.33<sup>b</sup> | 11.67±0.00<sup>a</sup> | 26.00±2.00<sup>d</sup> | - |
| *P. aeruginosa* | 16.67±0.33<sup>d</sup> | 14.67±0.33<sup>c</sup> | 11.67±0.33<sup>b</sup> | 8.33±0.33<sup>a</sup> | 0.00±0.00<sup>a</sup> | - |

Values are presented as mean ± SE of triplicates. Values with the same superscript letter(s) along the same rows are not significantly different ( $P < 0.05$ ) according to Tukey's Honestly Significant Difference
KEY: C (+): Positive Control (CiproTab - 2mg/ml), C (-): Negative Control – n-Hexane.

The observed antimicrobial activity of the extracts of *S. mombin* in this study aligns with existing research on the plant’s medicinal properties. Previous studies have reported the presence of various bioactive compounds in *S. mombin*, including alkaloids, flavonoids, and phenols, which have been associated with antimicrobial activity [26]. Additionally, the effectiveness of *S. mombin* against antibiotic-resistant bacteria corroborates findings from other studies that have investigated the antimicrobial potential of plant extracts against multidrug-resistant pathogens [27].

The results presented in Tables 6, 7, and 8 highlight the antibacterial effectiveness of different extracts of *S. mombin* against multiple antibiotic-resistant bacterial isolates obtained from urine samples. These findings contribute to the existing research on the antimicrobial properties of *S. mombin* and support its potential use as an alternative therapeutic option for treating antibiotic-resistant urinary tract infections [28].

**Table 6:** Antibacterial activity of aqueous extract on isolates from urine samples.

| Isolates | 100 | 50 | 25 | 12.5 | C(+) | C(-) |
| --- | --- | --- | --- | --- | --- | --- |
|  | (mg/ml) |  |  |  |  |  |
| *E. coli* | 24.67±0.33<sup>d</sup> | 21.00±0.00<sup>c</sup> | 16.00±0.00<sup>b</sup> | 10.33±0.33<sup>a</sup> | 29.00±1.00<sup>d</sup> | - |
| *K. pneumoniae* | 27.33±0.33<sup>d</sup> | 24.33±0.33<sup>c</sup> | 18.33±0.33<sup>b</sup> | 12.67±0.33<sup>a</sup> | 31.00±1.00<sup>d</sup> | - |
| *P. aeruginosa* | 18.67±0.33<sup>d</sup> | 15.67±0.33<sup>c</sup> | 11.67±0.33<sup>b</sup> | 0.00±0.00<sup>a</sup> | 24.00±1.00<sup>d</sup> | - |
| *P. mirabilis* | 24.67±0.33<sup>d</sup> | 22.33±0.33<sup>c</sup> | 18.00±0.00<sup>b</sup> | 11.00±0.00<sup>a</sup> | 34.00±1.00<sup>d</sup> | - |
| *S. aureus* | 23.67±0.33<sup>d</sup> | 15.67±0.33<sup>c</sup> | 10.67±0.33<sup>b</sup> | 9.00±0.00<sup>a</sup> | 22.00±1.00<sup>d</sup> | - |
| *S. saprophyticus* | 21.33±0.33<sup>d</sup> | 17.67±0.33<sup>c</sup> | 11.67±0.33<sup>b</sup> | 9.00±0.00<sup>a</sup> | 28.33±0.33<sup>d</sup> | - |
| *T. vaginalis* | 21.67±0.33<sup>d</sup> | 16.67±0.33<sup>c</sup> | 12.33±0.33<sup>b</sup> | 9.00±0.00<sup>a</sup> | 33.33±0.33<sup>d</sup> | - |

Values are presented as mean ± SE of triplicates. Values with the same superscript letter(s) along the same rows are not significantly different ( $P < 0.05$ ) according to Tukey's Honestly Significant Difference
KEY: C (+): Positive Control (CiproTab - 2mg/ml), C (-): Negative Control - Water.

**Table 7:** Antibacterial activity of methanol extract on isolates from urine samples.

| Isolates | 100 | 50 | 25 | 12.5 | C(+) | C(-) |
| --- | --- | --- | --- | --- | --- | --- |
|  | (mg/ml) |  |  |  |  |  |
| *E. coli* | 22.00±0.00<sup>d</sup> | 18.00±0.00<sup>c</sup> | 13.00±0.00<sup>b</sup> | 7.00±0.00<sup>a</sup> | 29.00±1.00<sup>d</sup> | - |
| *K. pneumoniae* | 25.00±0.00<sup>d</sup> | 21.00±0.00<sup>c</sup> | 18.00±0.00<sup>b</sup> | 14.00±0.00<sup>a</sup> | 31.00±1.00<sup>d</sup> | - |
| *P. aeruginosa* | 17.00±0.00<sup>d</sup> | 15.00±0.00<sup>c</sup> | 12.00±0.00<sup>b</sup> | 0.00±0.00<sup>a</sup> | 24.00±1.00<sup>d</sup> | - |
| *P. mirabilis* | 20.00±0.00<sup>d</sup> | 17.67±0.33<sup>c</sup> | 13.00±0.00<sup>b</sup> | 0.00±0.00<sup>a</sup> | 34.00±1.00<sup>d</sup> | - |
| *S. aureus* | 20.67±0.33<sup>d</sup> | 18.00±0.00<sup>c</sup> | 14.67±0.33<sup>b</sup> | 11.00±0.00<sup>a</sup> | 22.00±1.00<sup>d</sup> | - |
| *S. saprophyticus* | 18.33±0.33<sup>d</sup> | 15.67±0.33<sup>c</sup> | 13.00±0.00<sup>b</sup> | 9.00±0.00<sup>a</sup> | 28.33±0.33<sup>d</sup> | - |
| *T. vaginalis* | 20.33±0.33<sup>d</sup> | 15.67±0.33<sup>c</sup> | 13.00±0.00<sup>b</sup> | 9.00±0.00<sup>a</sup> | 33.33±0.33<sup>d</sup> | - |

Values are presented as mean ± SE of triplicates. Values with the same superscript letter(s) along the same rows are not significantly different ( $P<0.05$ ) according to Tukey's Honestly Significant Difference
KEY: C (+): Positive Control (CiproTab - 2mg/ml), C (-): Negative Control - Methanol.

**Table 8:** Antibacterial activity of n-Hexane extract on isolates from urine samples.

| Isolates | 100 | 50 | 25 | 12.5 | C(+) | C(-) |
| --- | --- | --- | --- | --- | --- | --- |
|  | (mg/ml) |  |  |  |  |  |
| *E. coli* | 18.00±0.00<sup>d</sup> | 15.00±0.00<sup>c</sup> | 13.00±0.00<sup>b</sup> | 7.00±0.00<sup>a</sup> | 29.00±1.00<sup>d</sup> | - |
| *K. pneumoniae* | 19.00±0.00<sup>d</sup> | 15.00±0.00<sup>c</sup> | 11.00±0.00<sup>b</sup> | 9.00±0.00<sup>a</sup> | 31.00±1.00<sup>d</sup> | - |
| *P. aeruginosa* | 17.00±0.00<sup>d</sup> | 14.00±0.00<sup>c</sup> | 12.00±0.00<sup>b</sup> | 8.00±0.00<sup>a</sup> | 24.00±1.00<sup>d</sup> | - |
| *P. mirabilis* | 19.00±0.00<sup>d</sup> | 17.00±0.00<sup>c</sup> | 14.00±0.00<sup>b</sup> | 9.00±0.00<sup>a</sup> | 34.00±1.00<sup>d</sup> | - |
| *S. aureus* | 16.00±0.00<sup>d</sup> | 14.00±0.00<sup>c</sup> | 11.00±0.33<sup>b</sup> | 0.00±0.00<sup>a</sup> | 22.00±1.00<sup>d</sup> | - |
| *S. saprophyticus* | 18.00±0.00<sup>d</sup> | 16.00±0.00<sup>c</sup> | 13.00±0.00<sup>b</sup> | 0.00±0.00<sup>a</sup> | 28.33±0.33<sup>d</sup> | - |
| *T. vaginalis* | 15.00±0.00<sup>d</sup> | 11.00±0.00<sup>c</sup> | 9.00±0.00<sup>b</sup> | 0.00±0.00<sup>a</sup> | 33.33±0.33<sup>d</sup> | - |

Values are presented as mean ± SE of triplicates. Values with the same superscript letter(s) along the same rows are not significantly different ( $P<0.05$ ) according to Tukey's Honestly Significant Difference
KEY: C (+): Positive Control (CiproTab - 2mg/ml), C (-): Negative Control - n-Hexane.

The study reveals the antibacterial properties of *S. mombin* extracts against various antibiotic-resistant bacteria from urine samples. The aqueous extract showed the highest inhibition zone against *K. pneumoniae* (27.33±0.33d), while the methanolic extract showed the highest inhibition zone against all isolates, with zone diameters ranging from 17.00±0.00 to 25.00±0.00 at a concentration of 100 mg/ml. The n-Hexane extract also showed the highest inhibition zone against all isolates, with smaller zones of inhibition than the positive control. These findings support the potential of *S. mombin* as an alternative therapeutic option for treating antibiotic-resistant urinary tract infections. The results support the potential of *S. mombin* as a potential therapeutic option.

The findings of this study are consistent with previous research on the antimicrobial properties of *S. mombin* extracts. Other studies have reported the presence of bioactive compounds in *S. mombin*, such as flavonoids, tannins, and alkaloids, which are known for their antimicrobial activities [27]. Furthermore, studies investigating the antimicrobial potential of *S. mombin* extracts against antibiotic-resistant bacteria have shown promising results [7]. These findings support the notion that *S. mombin* extracts may serve as valuable resources for the development of new therapeutic agents against antibiotic-resistant urinary tract infections.

The higher effectiveness of the aqueous extract compared to the methanol and n-hexane extracts suggests that the water-soluble components of *S. mombin* may play a significant role in its antimicrobial activity. This finding aligns with previous studies that have reported the antimicrobial properties of aqueous extracts of *S. mombin* against various bacterial strains [29]. The presence of bioactive compounds such as alkaloids, flavonoids, phenols, and tannins in the aqueous extract, as identified in previous phytochemical analyses [30], may contribute to its observed antibacterial activity. The observed lower zones of inhibition for the n-hexane extract suggest that the non-polar compounds present in *S. mombin* may have limited effectiveness against the tested isolates. This aligns with the work of Trusheva *et al*. (2007), [31] who have reported that the varying antimicrobial activities of different solvent extracts from *S. mombin*, indicating that the choice of solvent can significantly influence the extraction of bioactive compounds with antimicrobial properties. In contrast, the aqueous extract of *S. mombin* showed the highest effectiveness against tested isolates, with larger zone diameters of inhibition. This suggests the extract contains water-soluble bioactive compounds, such as alkaloids, flavonoids, phenols, and tannins, which have antibacterial properties [30].

Table 9 presents MIC and MBC values of *S. mombin* extracts against wound samples, enhancing research on antimicrobial potential and determining effective concentrations for specific bacterial strains. The extracts showed a MIC value of 50 mg/ml against *P. aeruginosa*, with a MBC value of 100mg/ml, suggesting a higher concentration is needed for bactericidal effects, aligning with previous studies on *S. mombin* extracts [32]. *S. mombin* extracts showed lower MIC and MBC values against *S. epidermidis* compared to *P. aeruginosa*, indicating lower antibiotic resistance. These results suggest lower concentrations are needed for effective inhibitory and bactericidal effects. The study found that *S. mombin* extracts have moderate antimicrobial activity against *S. aureus* strains, with a MIC value of 25mg/ml for aqueous and methanol extracts, and 50mg/ml for n-hexane extract. Table 10 provides insights into the MIC and MBC values of *S. mombin* extracts against isolates derived from urine samples, including *E. coli, S. aureus, S. saprophyticus, T. vaginalis, K. pneumoniae*, P. mirabilis, and *P. aeruginosa*. The MIC and MBC values ranged from 25 mg/ml to 100mg/ml, indicating varying levels of susceptibility among the tested isolates. The range of MIC and MBC values suggests variations in the effectiveness of the extracts against different pathogens. These findings are consistent with previous studies that have reported variable susceptibility patterns of bacterial isolates to *S. mombin* extracts [32, 33].

**Table 9:** MIC and MBC values of isolates from wound samples.

| Isolates | MIC |  |  | MBC |  |  |
| --- | --- | --- | --- | --- | --- | --- |
|  | Aqueous | N-hexane | Methanol<br>(mg/ml) | Aqueous | N-hexane | Methanol |
| *S. aureus* | 25 | 50 | 25 | 50 | 100 | 50 |
| *S. epidermidis* | 12.5 | 12.5 | 12.5 | 25 | 25 | 25 |
| *P. aeruginosa* | 50 | 50 | 50 | 100 | 100 | 100 |

**Table 10:** MIC and MBC values of isolate from urine samples.

| Isolates | MIC |  |  | MBC |  |  |
| --- | --- | --- | --- | --- | --- | --- |
|  | Aqueous | N-hexane | Methanol<br>(mg/ml) | Aqueous | N-hexane | Methanol |
| *E. coli* | 50 | 50 | 50 | 100 | 100 | 100 |
| *K. pneumoniae* | 25 | 25 | 25 | 50 | 50 | 50 |
| *P. aeruginosa* | 25 | 50 | 25 | 50 | 100 | 50 |
| *P. mirabilis* | 25 | 25 | 25 | 50 | 50 | 50 |
| *S. aureus* | 50 | 50 | 50 | 100 | 100 | 100 |
| *S. saprophyticus* | 25 | 25 | 25 | 50 | 50 | 50 |
| *T. vaginalis* | 50 | 50 | 50 | 100 | 100 | 100 |

**Table 11:** Qualitative Phytochemical Profile of *S. mombin*.

| Phytochemicals | Water | Methanol | n-Hexane |
| --- | --- | --- | --- |
| Alkaloids | + | + | + |
| Anthraquinones | + | - | - |
| Cardiac Glycosides | + | + | - |
| Flavonoids | + | + | + |
| Phenols | + | + | + |
| Saponins | + | + | - |
| Steroids | + | - | + |
| Tannins | + | + | + |

Key- +:Positive, - :Negative

The qualitative analysis of *S. mombin* extracts revealed various phytochemical classes, including alkaloids, anthraquinones, cardiac glycosides, flavonoids, phenols, saponins, steroids, and tannins, which are known for their antimicrobial and antioxidant properties against antibiotic-resistant bacterial isolates. These compounds can contribute to the observed antimicrobial effects of *S. mombin* extracts against antibiotic-resistant bacterial isolates [30]. Anthraquinones were not detected in methanol and n-hexane extracts, suggesting they may not be the main antimicrobial constituents, possibly due to the extraction process and solvents. Saponins were also detected in the aqueous and methanol extracts but were not found in the n-hexane extract. Saponins are known for their diverse biological activities, including antimicrobial, anti-inflammatory, and anticancer properties [33]. The presence of saponins in *S. mombin* extracts further supports their potential as antimicrobial agents against multidrug-resistant bacterial isolates. The comprehensive analysis of bacterial isolates obtained from wound and urine samples, as well as the evaluation of *S. mombin* extracts, provided valuable insights into the prevalence of antibiotic-resistant bacteria and the potential antibacterial properties of the plant.

## Conclusion

Antibiotic-resistant bacteria pose a global health challenge, necessitating novel therapeutic approaches. *S. mombin* plant extracts offer potential for developing new antimicrobial agents, highlighting the importance of exploring natural products and their bioactive compounds to combat antibiotic resistance and improve public health.

## Recommendation

Further research is needed to identify bioactive compounds responsible for antimicrobial activity, understand their mechanisms, and conduct in vivo experiments and clinical trials for potential therapeutic applications.

## Contributions to knowledge

The research highlights the alarming prevalence of antibiotic-resistant bacteria in wound and urine samples within the study area and explores the potential of *S. mombin* plant extracts as a natural, effective treatment option for these infections.

### Funding

This research was not funded by any organization

### Ethical Approval

Obtained from the Ethics and Research Section of Ondo State Ministry of Health.

### Authors’ contributions

This work was carried out in collaboration between all authors. Author AFO designed the study, performed the statistical analysis, wrote the protocol and wrote the first draft of the manuscript. Author MKO and AOO managed the analyses of the study and were in charge of direction and planning.

## Acknowledgements

Special thanks to the Department of Microbiology, The Federal University of Technology Akure (FUTA) for making this research possible.

## Abbreviations

CAZ: Ceftazidime
CRX: Cefuroxime
GEN: Gentamycin
CTR: Ceftriazone
ERY: Erythromycin
CXC: Cloxacillin
OFL: Ofloxacin
AUG: Augmentin
CXM: Cefixime
NIT: Nitrofurantoin
CPR: Cefpiroma
MSA: Manitol salt agar
CLED: Cysteine lactose electrolyte-deficient
mg/ml: milligram per ml.

## REFERENCES

1. World Health Organization: WHO. Antimicrobial resistance [Internet]. 2023. Available from: https://www.who.int/news-room/fact-sheets/detail/antimicrobial-resistance (Accessed 27th March 2023)

2. Dadgostar P. Antimicrobial resistance: implications and costs. Infection and drug resistance. 2019 Dec 20:3903–10.

3. Laxminarayan R, Duse A, Wattal C, Zaidi AK, Wertheim HF, Sumpradit N, et al. Antibiotic resistance—the need for global solutions. The Lancet infectious diseases. 2013 Dec 1;13(12):1057–98.

4. O’neill JI. Antimicrobial resistance: tackling a crisis for the health and wealth of nations. Rev. Antimicrob. Resist.. 2014.

5. Chaachouay N, Zidane L. Plant-Derived Natural Products: A Source for Drug Discovery and Development. Drugs and Drug Candidates. 2024;3(1):184–207.

6. Mahajan GB, Balachandran L. Antibacterial agents from actinomycetes-a review. Frontiers in Bioscience-Elite. 2012 Jan 1;4(1):240–53.

7. Ogunro OB, Oyeyinka BO, Gyebi GA, Batiha GE. Nutritional benefits, ethnomedicinal uses, phytochemistry, pharmacological properties and toxicity of Spondias mombin Linn: a comprehensive review. Journal of Pharmacy and Pharmacology. 2023 Feb 1;75(2):162–226.

8. Ojewole JA, Adewole SO, Olayiwola G. Hypoglycaemic and hypotensive effects of Momordica charantia Linn (Cucurbitaceae) whole-plant aqueous extract in rats: Cardiovascular topics. Cardiovascular Journal of South Africa. 2006 Sep 1;17(5):227–32.

9. Nigussie D, Davey G, Legesse BA, Fekadu A, Makonnen E. Antibacterial activity of methanol extracts of the leaves of three medicinal plants against selected bacteria isolated from wounds of lymphoedema patients. BMC Complementary Medicine and Therapies. 2021 Dec;21:1–0.

10. Pernille H, Lars B, Marjukka M, Volkert S, Anne H. Sampling of urine for diagnosing urinary tract infection in general practice–First-void or mid-stream urine?. Scandinavian journal of primary health care. 2019 Jan 2;37(1):113–9.

11. Oluyele O, Oladunmoye M. Antibiotics Susceptibility Patterns and Plasmid Profile of Staphylococcus aureus Isolated from Patients with Wound Infections Attending Four Hospitals in Akure, Ondo State. Journal of Advances in Microbiology. 2017:3(4):1–8.

12. Shoaib M, Muzammil I, Hammad M, Bhutta ZA, Yaseen I. A mini-review on commonly used biochemical tests for identification of bacteria. International Journal of Research Publications. 2020 Jun 14;54(1):1–7.

13. Cheesbrough M. District laboratory practice in tropical countries, part 2. Cambridge university press; 2005.

14. Okiti AF, Osuntokun OT. Antimicrobial, phytochemical analysis and molecular docking (In-silico Approach) of Tithonia diversifolia (Hemsl.) A. Gray and Jatropha gossypiifolia L on selected clinical and multi-drug resistant isolates. Journal of Advances in Microbiology. 2020;20(6):1–8.

15. Ogbonnia SO, Odimegwu JI, Enwuru VN. Evaluation of hypoglycaemic and hypolipidaemic effects of aqueous ethanolic extracts of Treculia africana Decne and Bryophyllum pinnatum,/i> Lam. and their mixture on streptozotocin (STZ)-induced diabetic rats. African Journal of Biotechnology. 2008;7(15).

16. Rankin DI, Coyle BM. Manual of antimicrobial susceptibility testing. American Society for Microbiology. 2005;53:62.

17. National Committee for Clinical Laboratory Standards, Barry AL. Methods for determining bactericidal activity of antimicrobial agents: approved guideline. Wayne, PA: National Committee for Clinical Laboratory Standards; 1999 Sep.

18. Mallikharjuna PB, Rajanna LN, Seetharam YN, Sharanabasappa GK. Phytochemical studies of Strychnos potatorum Lf-A medicinal plant. Journal of chemistry. 2007;4(4):510–8.

19. Kumar GS, Jayaveera KN, Kumar CK, Sanjay UP, Swamy BM, Kumar DV. Antimicrobial effects of Indian medicinal plants against acne-inducing bacteria. Tropical journal of pharmaceutical research. 2007 Jul 31;6(2):717–23.

20. Onwukaeme DN, Ikuegbvweha TB, Asonye CC. Evaluation of phytochemical constituents, antibacterial activities and effect of exudate of Pycanthus Angolensis Weld Warb (Myristicaceae) on corneal ulcers in rabbits. Tropical Journal of Pharmaceutical Research. 2007;6(2):725–30.

21. Hazra KM, Roy RN, Sen SK, Laskar S. Isolation of antibacterial pentahydroxy flavones from the seeds of Mimusops elengi Linn. African journal of Biotechnology. 2007;6(12).

22. Parekh J, Chanda S. In vitro antimicrobial activity and phytochemical analysis of some Indian medicinal plants. Turkish journal of biology. 2007;31(1):53–8.

23. Liu G, Qin M. Analysis of the distribution and antibiotic resistance of pathogens causing infections in hospitals from 2017 to 2019. Evidence-Based Complementary and Alternative Medicine. 2022;2022(1):3512582.

24. Ferreira RL, Da Silva BC, Rezende GS, Nakamura-Silva R, Pitondo-Silva A, Campanini EB, et al. High prevalence of multidrug-resistant Klebsiella pneumoniae harboring several virulence and β-lactamase encoding genes in a Brazilian intensive care unit. Frontiers in microbiology. 2019 Jan 22;9:3198.

25. Li Z, Peng C, Zhang G, Shen Y, Zhang Y, Liu C, et al. Prevalence and characteristics of multidrug-resistant Proteus mirabilis from broiler farms in Shandong Province, China. Poultry science. 2022 Apr 1;101(4):101710.

26. Gopu C, Chirumamilla P, Daravath SB, Vankudoth S, Taduri S. GC-MS analysis of bioactive compounds in the plant parts of methanolic extracts of Momordica cymbalaria Fenzl. J. Med. Plants Stud. 2021;9(3):209–18.

27. Osuntokun OT. Evaluation of Inhibitory Zone Diameter (IZD) of crude Spondias mombin (Linn.) extracts (root, leaf, and stem bark) against thirty infectious clinical and environmental isolates. J Bacteriol Infec Dis. 2018; 2 (1): 8. 2018;16.

28. Bossou AF, Dokpè S, Atindehou M, Sophie G, Bogninou R, Koudoro Y, et al. Evaluation of antiradical and antibacterial activities of hydroethanolic extract of Spondias mombin Leaves from Benin. IOSR J Pharm. 2020;10:11–6.

29. de Lima IP, Alves RA, Mayer JD, Costa MR, de Mendonça AK, de Lima EL, et al. Antimicrobial activity of Spondias mombin L. aqueous and hydroethanolic extracts on Enterococcus faecalis and Pseudomonas aeruginosa-an in vitro study. Research, Society and Development. 2021 Jan 28;10(1):e50710111949..

30. Saha S. Phytochemical screening and comparative study of antioxidative properties of the fruits and leaves of Spondias mombin in Bangladesh. Journal of Pharmacognosy and Phytochemistry. 2019;8(2):379–83.

31. Trusheva B, Trunkova D, Bankova V. Different extraction methods of biologically active components from propolis: a preliminary study. Chemistry Central Journal. 2007 Dec;1:1–4.

32. Temitope OO, Ogunmodede AF, Fasusi OA, Thonda AO, Odufunwa AE. Synergistic antibacterial and antifungal activities of Spondias mombin extracts and conventional antibiotic and antifungal agents on selected clinical microorganisms. Sch. J. Appl. Med. Sci. 2017;5:307–18.

33. Barbosa AD. An overview on the biological and pharmacological activities of saponins. Int J Pharm Pharm Sci. 2014;6(8):47–50.

